# Comparative transcriptome analysis of human and murine choroidal neovascularization identifies fibroblast growth factor inducible-14 as phylogenetically conserved mediator of neovascular age-related macular degeneration

**DOI:** 10.1101/2021.05.10.443381

**Authors:** Julian Wolf, Anja Schlecht, Dennis-Dominik Rosmus, Stefaniya Boneva, Hansjürgen Agostini, Günther Schlunck, Peter Wieghofer, Clemens Lange

## Abstract

**Background:** Visual outcome of patients with neovascular age-related macular degeneration has significantly improved during the last years following the introduction of anti-vascular endothelial growth factor (VEGF) therapy. However, about one third of patients show persistent exudation and decreasing visual acuity despite recurrent anti-VEGF treatment, which implies a role of other, still unknown proangiogenic mediators.

**Methods:** The present study applied transcriptional profiling of human and mouse (C57BL/6J wildtype) choroidal neovascularization (CNV) membranes each with reference to healthy control tissue to identify yet unrecognized mediators of CNV formation. Key factors were further investigated by immunohistochemistry as well as by intravitreal inhibition experiments and multiplex protein assays in the laser-induced CNV mouse model.

**Results:** Transcriptional profiles of CNV membranes were characterized by enhanced activation of blood vessel development, cytoskeletal organization, and cytokine production, with angiogenesis and wound healing processes predominating in humans and activation of immune processes in mice. Besides several species-specific factors, 95 phylogenetically conserved CNV-associated genes were detected, among which fibroblast growth factor inducible-14 (FN14), a member of the tumor necrosis factor (TNF) receptor family, was identified as a key player of CNV formation. Blocking the pathway by intravitreal injection of a FN14 decoy receptor modulated the cytokine profile - most notably IL-6 - and led to a significant reduction of CNV size *in vivo*.

**Conclusions:** This study characterizes the transcriptome of human and mouse CNV membranes in an unprejudiced manner and identifies FN14 as a phylogenetically conserved mediator of CNV formation and a promising new therapeutic target for neovascular AMD.

**Funding:** This study was funded by the Helmut-Ecker-Stiftung and the Volker-Homann-Stiftung.

## Introduction

Age-related macular degeneration (AMD) is the most common cause of permanent blindness in elderly people in developed countries. Its worldwide prevalence is estimated to increase from 200 million in 2020 to 300 million by 2040 as a result of population ageing (Wong et al., 2014). Worldwide, about 40 million people suffer from the neovascular form of AMD (nAMD) (Wong et al., 2014), which is characterized by the formation of pathological choroidal neovascularisation (CNV) leading to edema, hemorrhage, scarring and ultimately irreversible impairment of central vision (Mitchell et al., 2018). An important proangiogenic mediator in this process is the vascular endothelial growth factor (VEGF), which increases vascular permeability and promotes the development of CNV. Within the last years, anti-VEGF therapy has significantly improved visual outcomes in nAMD (Bressler et al., 2011; Mitchell et al., 2018). However, about one third of patients with nAMD show persistent exudation and a slow decrease in visual acuity despite intensive anti-VEGF therapy (Wecker et al., 2019; Yang et al., 2016). This illustrates the complexity of the disease and indicates a role of other proangiogenic mediators in the development and progression of CNV in nAMD.

So far, the laser-induced CNV mouse model was extensively used to mimic the disease *in vivo* allowing the identification and evaluation of novel mediators of CNV development (Bucher et al., 2020; Lange et al., 2016; Schlecht et al., 2020b; Wang et al., 2013). However, an unbiased comparison of the transcriptional landscapes between murine and human CNV is still lacking but of significant importance for the interpretation of the mouse model and the translation of preclinical findings into novel clinical treatment regimens. To address the urgent need for alternative therapeutic approaches, the present study characterizes the transcriptome of human and mouse CNV membranes each with reference to healthy control tissue, analyses the involved biological processes, and compares differentially expressed genes in order to identify conserved mediators of CNV formation. Following this approach, the study identifies fibroblast growth factor inducible-14 (FN14) as a novel key factor of CNV formation, investigates its influence on the cytokine milieu of CNV, as well as its antiangiogenic effect *in vivo*.

## Methods

### Patients

CNV membranes were obtained from four patients with neovascular AMD undergoing the meanwhile obsolete procedure of subretinal CNV extraction (Bressler et al., 2000) between 1992 and 1999. Only patients with treatment-naive classical CNV associated with typical AMD changes such as drusen and RPE alterations were included in the study. Patients with concomitant diseases, such as myopia or central serous chorioretinopathy, were excluded from the study. Following a 20-gauge vitrectomy, a retinotomy was performed temporally to the macula and the CNV membrane was surgically extracted using a flexed forceps. Four age-matched RPE-choroidal samples from the macular region of four enucleated eyes suffering from ciliary body melanoma served as controls. For protein analysis, undiluted vitreous samples were obtained from further patients with nAMD (n = 6) and control patients (n = 6) who underwent vitrectomy for subretinal bleeding or macular pucker, respectively. Ethics approval was granted from local Ethics Committees.

### Mice

Mice used in this study were C57BL/6J wildtype mice purchased from Charles River. All animal experiments were approved by the local authority (Regierungspräsidium Freiburg, Germany) and were performed in accordance with the respective national, federal and institutional regulations. All animals were tested negatively for the Rd8 (*Crb1*) mutation.

### Laser-induced choroidal neovascularization model

Six to eight weeks old C57BL/6J wildtype mice were anesthetized with intraperitoneal injection of a mixture of ketamine (100 µg/g) and xylazine (12 µg/g). Pupils were dilated with a combination of tropicamide (5 mg/ml) and neosynephrine-POS (50 mg/ml). Lubricating gel was administered to protect the eye and to maintain corneal hydration. The eye was carefully placed against a cover glass to flatten the curved corneal surface. Laser burns at equal distance from the optic disc were induced by an Argon laser (VISULAS 532s, ZEISS) with a wavelength of 532 nm, a power of 150 mW, a diameter of 100 µm and duration of 100 ms (Buhler et al., 2013). Only burns that produced a bubble as a sign for Bruch’s membrane rupture and without concomitant bleeding were included in the study (Tobe et al., 1998). After laser treatment, mice were placed on a prewarmed warming plate at 35 °C until they recovered from anesthesia.

### Intravitreal injection of a FN14 decoy receptor and quantification of CNV lesion size

Laser-induced choroidal neovascularization (CNV) was induced at day 0 (d0) in C57BL/6J wildtype mice as described above. At d1 mice received an intravitreal injection of 4 µg FN14 decoy receptor (1610-TW, R&D Systems) dissolved in 1 µl PBS in one eye and the same volume of PBS in the other eye. Injections were performed with a Nanofil 10 µl syringe using a Nanofil 34 G needle (World Precision Tools, Sarasota, USA) under microscopic visual control. At d7, fundus photography and fluorescein angiography (ALCON Fluorescein 10 %, diluted 1:20 with 0.9 % NaCl, intraperitoneal injection of 40 µL per 20 g mouse) were performed using a retinal imaging microscope (Micron IV, Phoenix Technology Group, Pleasanton, USA). To quantify CNV size, RPE/choroidal flatmounts were dissected and stained (see section below) and fluorescence images (647 nm, COL4) of the entire RPE/choroid were taken using a Hamamatsu NanoZoomer S60 (Hamamatsu Photonics, Herrsching, Germany). Spot sizes were determined by a masked investigator. Confluent lesions and animals with vitreal or subretinal hemorrhage were excluded from further analysis. Each inhibition experiment was independently performed twice.

### Sample preparation for RNA sequencing

An illustration of the experimental setup for transcriptional characterization of human and mouse CNV is shown in Figure 1A. For transcriptomic characterization of human CNV membranes, RNA sequencing data generated and recently published by our group (Schlecht et al., 2020a) were reanalyzed as described below. FFPE processing was performed immediately after resection according to routine protocols, as described previously (Lange et al., 2018; Schlecht et al., 2020a). Following routine histological staining, each specimen’s histological diagnosis was made by two experienced ophthalmic pathologists. Fifteen FFPE sections of 4 µm thickness from each CNV membrane and each central RPE-choroid complex were stored in tubes prior to RNA extraction, as previously described (Schlecht et al., 2020a).

**Figure 1:**
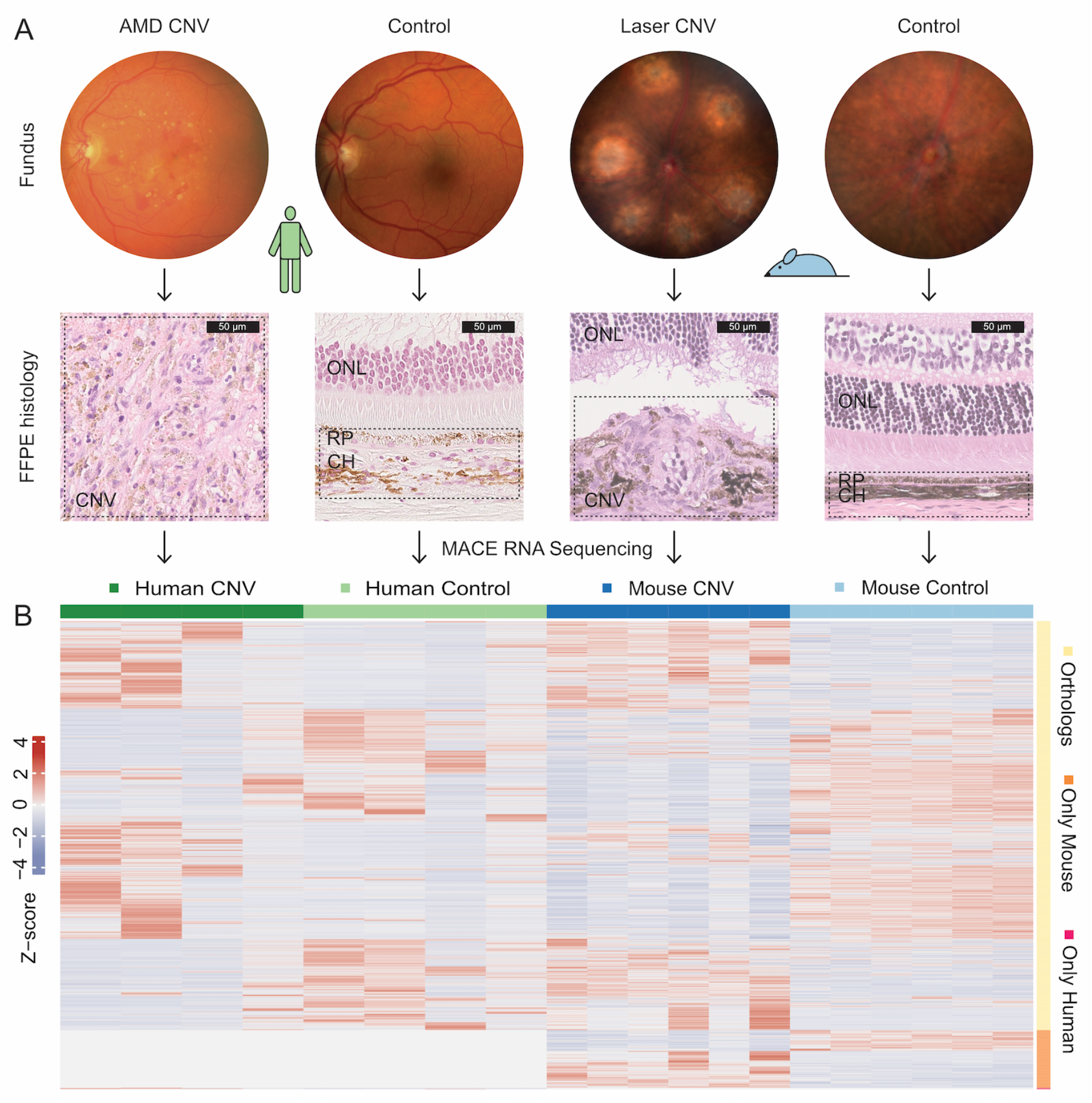
Experimental setup for transcriptional characterization of human and mouse CNV. (A) Representative fundus photography (upper panel) and hematoxylin eosine stainings (lower panel) of human and mouse CNVs and controls. A representative tissue area used for RNA sequencing is indicated by dashed boxes (lower panel). Abbreviations: AMD: age-related macular degeneration, CH: choroid, CNV: choroidal neovascularization, ONL: outer nuclear layer, RP: retinal pigment epithelium. (B) Supervised heatmap of differentially expressed genes (DEG) between CNV and control samples of both species. Each column represents one sample and each row one DEG. Among the 4926 DEGs there was a one-to-one ortholog of the other species for 4305 genes (87.4 %) (no orthologous gene for 606 (12.3 %) mouse and 15 (0.3 %) human DEGs, see Heatmap annotation to the right). The z-score represents a gene’s expression in relation to its mean expression by standard deviation units (red: upregulation, blue: downregulation).

For transcriptome analysis of mouse CNV membranes, CNV was induced in six C57BL/6J wildtype mice with six laser spots per eye at d0, as described above. Six mice without laser treatment served as controls. At d7, fundus photography was performed (Micron IV, Phoenix Technology Group, Pleasanton, USA) and mice were sacrificed by cervical dislocation. After enucleation, the eyes were immediately fixed in 4 % formaldehyde for 2 h at room temperature (RT) as previously described (Lange et al., 2018; Schlecht et al., 2020a) followed by extensive washing in PBS. Subsequently, central RPE and choroid were dissected in PBS. FFPE RPE-choroid samples from both eyes of one animal were pooled and stored in tubes prior to RNA extraction. In addition, routine hematoxylin and eosin staining was performed on 4 µm sections of FFPE processed eyes of the laser and control group. FFPE processing was kept identical for human and mouse samples to ensure methodological comparability.

### RNA isolation

RNA isolation from FFPE specimens was carried out as previously described (Lange et al., 2018; Schlecht et al., 2020a). Briefly, total RNA was extracted from FFPE samples using the Quick-RNA FFPE Kit (Zymo Research, Irvine, California). Following DNAse I digestion using the Baseline-ZERO Kit (Epicentre, Madison, Wisconsin), the RNA concentration was quantified using the Qubit RNA HS Assay Kit on a Qubit Fluorometer (Life Technologies, Carlsbad, California). RNA quality was determined via the RNA Pico Sensitivity Assay on a LabChip GXII Touch (PerkinElmer, Waltham, Massachusetts).

### MACE RNA-Sequencing

RNA sequencing was performed using massive analysis of cDNA ends (MACE), a 3’ RNA sequencing method, as previously described (Schlecht et al., 2020a; Wolf et al., 2020). We recently demonstrated that MACE allows sequencing of FFPE samples with high accuracy independent of storage time (Boneva et al., 2020). Briefly, 20 barcoded libraries comprising unique molecule identifiers (six mouse CNV membranes, six mouse RPE-choroidal control samples, four human CNV membranes and four human RPE-choroidal control tissues) were sequenced on the NextSeq 500 (Illumina) with 1 × 75 bp. PCR bias was removed using unique molecular identifiers.

### Bioinformatics

Sequencing data (fastq-files) were uploaded to the Galaxy web platform (usegalaxy.eu) (Afgan et al., 2018), as previously described (Boeck et al., 2020). Quality control was performed with FastQC Galaxy Version 0.72 (http://www.bioinformatics.babraham.ac.uk/projects/fastqc/ last access on 08/04/2019). Reads were mapped to the human (hg38) or mouse (mm10) Galaxy built-in reference genome with RNA STAR Galaxy Version 2.6.0b-2 (Dobin et al., 2013) using Gencode annotation files (human: Gencode 31, release June 2019, https://www.gencodegenes.org/human/releases.html, mouse: Gencode M22, release June 2019, https://www.gencodegenes.org/mouse/releases.html). Reads mapped to the human or mouse reference genome were counted using featureCounts (Galaxy Version 1.6.4) (Liao et al., 2014) using the aforementioned annotation files. The outputs of featureCounts were imported to RStudio (Version 1.2.1335, R Version 3.5.3). Gene symbols and gene types were determined based on the ENSEMBL database (human genes, GRCh38.p12, download on 08/31/2019, mouse genes GRCm38.p6, download on 08/25/2019) (Zerbino et al., 2018). Orthologous genes were determined based on ENSEMBL one-to-one orthologs (download on 10/03/2019). After principle component analysis (Love et al., 2014), differential gene expression was analyzed using the R package DESeq2 1.22.2 with default parameters (Benjamini-Hochberg adjusted p-value) (Love et al., 2014). Transcripts with log2fold change (log2FC) > 0.5 or < -0.5 and adjusted p-value < 0.05 were considered as differentially expressed genes (DEG) (Schurch et al., 2016). Heatmaps were created with the R package ComplexHeatmap 1.20.0 (Gu et al., 2016). Gene enrichment analysis and its visualization with dotplots and cnetplots was done using the R package clusterProfiler 3.10.1 (Yu et al., 2012). Other data visualization was performed using the ggplot2 package (Wickham, 2016). The percentage of upregulated genes involved in disease-relevant biological processes was determined based on lists of genes associated with the GO-terms angiogenesis (GO:0001525), blood vessel development (GO:0001568) (combined to angiogenesis), immune response (GO:0006955) and response to wounding (GO:0009611) for human and mouse separately (download from http://geneontology.org (The Gene Ontology, 2019), on 10/21/2019). The sequencing data have been deposited to the Gene Expression Omnibus database under the accession number GSE163090.

### Immunohistochemistry

Mice were sacrificed by transcardial perfusion with phosphate-buffered saline (PBS) and 4 % paraformaldehyde (PFA) for 1 min each (10 ml/min, 4 °C). After enucleation, the eyes were immediately fixed in 4 % PFA for 1 h at 4 °C followed by extensive washing with PBS. Subsequently, flatmounts of retina and RPE/choroid were dissected in PBS. Flatmounts were then blocked and permeabilized with PBS containing 1 % bovine serum albumin (BSA) and 0.3 % Triton-X 100 overnight. Primary antibodies against Collagen IV (COL4, AB769, Merck Millipore, Billerica, MA, USA) and AIF1 (also known as IBA1, AB178846, Abcam, Cambridge, United Kingdom) in PBS containing 1 % BSA and 0.3 % Triton-X 100 were added over two nights at a dilution of 1:500 at 4 °C. After extensive washing with PBS containing 0.5 % BSA and 0.15 % Triton-X 100, secondary antibodies were added at a dilution of 1:500 (Alexa Fluor 647 donkey anti-goat (A21447), and Alexa Flour 568 donkey anti-rabbit (A10042), Thermo Fisher Scientific, Waltham, USA) overnight at 4 °C in the dark. After extensive washing with PBS, samples were mounted in Fluorescence Mounting Medium (Agilent Dako, Santa Clara, California, USA).

For cryosections, eyes were immediately fixed in 4 % PFA for 1 h at 4 °C followed by extensive washing in PBS. After corneotomy and lentectomy, eyes were dehydrated in 20 % sucrose overnight at 4 °C and embedded in Tissue-Tek O.C.T. Compound (Sakura Finetek Germany GmbH). Following extensive washing with PBS, 12 µm sections were blocked and permeabilized with PBS containing 1 % BSA and 0.3 % Triton-X 100 for 1 hour. Primary antibodies against FN14 (dilution: 1:500, ab109365, Abcam, Cambridge, United Kingdom), COL4 (dilution: 1:100, AB769, Merck Millipore, Billerica, MA, USA), AIF1 (dilution: 1:1000, ab178846, Abcam, Cambridge, United Kingdom) and rabbit monoclonal IgG isotype control (same concentration as FN14 antibody: 0.59 µg/µl, dilution: 1:500, ab172730, Abcam, Cambridge, United Kingdom) were incubated in PBS containing 1 % BSA and 0.3 % Triton-X 100 overnight at 4°C. For the negative control the primary antibody was omitted. Following extensive washing with PBS, secondary antibodies were incubated in PBS containing 1 % BSA and 0.3 % Triton-X 100 at a dilution of 1:500 (Alexa Fluor 488 donkey anti-rabbit (A21206) and 647 donkey anti-goat (A21447), Thermo Fisher Scientific, Waltham, USA) for 1 hour at RT in the dark. After extensive washing with PBS, nuclei were stained with 4′,6-Diamidin-2-phenylindol (DAPI) 1:1000 in PBS for 10 min at room temperature in the dark, washed again with PBS and mounted in Fluorescence Mounting Medium (Agilent Dako, Santa Clara, California, USA). Images were taken using a Leica SP8 confocal microscope (Leica, Wetzlar, Germany) equipped with a 20x NA 0.75 CS2 objective.

For Immunohistochemistry on human samples, human choroidal CNV-membranes from the Eye Center, University of Freiburg, and control FFPE samples from body donors at the Institute of Anatomy, Leipzig University, were used for immunohistochemistry. Eyes were enucleated in accordance with the consent of the body donors, which was secured by contract during lifetime, and no other data than age, sex, body weight and cause of death were disclosed. Eyes were fixed with 4 % PFA for 16 h at 4 °C and dissected under a binocular microscope. Paraffin embedding was performed using a standard protocol and 7 µm sections were obtained. Subsequently, paraffin sections were deparaffinized in accordance with a standard protocol and blocked with 2 % BSA and 2 % normal donkey serum (NDS) in PBS Triton-X 0.1 % for 60 min at RT. Primary antibodies were added at a dilution of 1:500 for FN14 (ab109365, Abcam, Cambridge, United Kingdom), 1:500 for AIF1 (234-013, Synaptic Systems) and 1:200 for Collagen IV (AB769, Millipore) in PBS containing 2 % BSA and 2 % NDS in PBS Triton-X 0.1 % overnight at 4 °C. Following extensive washing with 2 % BSA and 0.2 % NDS in PBS Triton-X 0.1 %, secondary antibodies were added at a dilution of 1:500 (Alexa Fluor^®^ 568 and Alexa Fluor^®^ 647, Thermo Fisher Scientific) in PBS Triton-X 0.1% at RT for 90 min in the dark. After washing at least three times with 2 % BSA and 0.2 % NDS in PBS Triton-X 0.1 %, slides were counterstained with DAPI 1:10000 for 10 min, washed three times with PBS followed by autofluorescence quenching with TrueBlack^®^ Lipofuscin Autofluorescence Quencher (Biotium) according to the manufacturer’s instructions. Slides were imaged using a confocal Fluoview FV1000 (Olympus) equipped with 20x 0.75 NA U Plan S Apo and 40x NA U Plan S objectives (Olympus).

### ELISA

FN14 is one of the main receptors of TWEAK (tumor necrosis factor related weak inducer of apoptosis). TWEAK concentration in human and murine CNV compared to the respective control tissue was determined by ELISA. For human CNV, undiluted vitreous samples were obtained from six patients with nAMD and six control patients who underwent vitrectomy for subretinal bleeding or macular pucker, respectively. Plasma samples were collected by venous puncture from the same patients at the time of surgery. The vitreous and blood samples were centrifuged for 20 min at 4 °C at 500 g and 15 min at 4 °C at 2000 g, respectively and were then transferred directly into sterile plastic tubes on ice. The samples were aliquoted and stored at -80 °C until analysis. Frozen vitreous and plasma samples were thawed and TWEAK levels were measured by using a human TWEAK ELISA kit (LS-F36324, LSBio) according to the manufacturer’s instructions. For mouse, CNV was induced in six C57BL/6J wildtype mice with six laser spots per eye at d0 as described above. Another six mice without CNV induction served as control. At d7, mice were sacrificed by cervical translocation. After enucleation, central RPE/choroid and retina were dissected and immediately frozen at -80 °C. Central RPE/choroid and retinae of both eyes of each animal were pooled for further analysis. Proteins were isolated from RPE/choroid as well as from retinal tissue of laser-treated and control mice using RIPA buffer (R0278, Sigma-Aldrich, St. Louis, Missouri, USA) containing protease (Complete Tablets Mini, 0463159001, Roche, Basel, Switzerland) and phosphatase inhibitors (Phosstop, 04906845001, Roche, Basel, Switzerland). Protein concentration was evaluated by a colorimetric assay (PierceTM BCA Protein Assay Kit, 23225, Thermo Fisher Scientific). A mouse TWEAK ELISA kit (LS-F32111, LSBio) was used to determine TWEAK concentrations in RPE/choroid and retinal samples according to the manufacturer’s instructions.

### Multiarray protein analysis

At d0 six laser spots were induced per eye in six mice, as described above. At d1 mice received an intravitreal injection of 4 µg FN14 decoy receptor (1610-TW, R&D Systems) solved in 1 µl PBS in one eye and the same volume of PBS in the other eye, as described above. Another five mice without CNV induction served as control. At d5, mice were sacrificed by cervical translocation. After enucleation, central RPE/choroid and retina were dissected and immediately frozen at -80 °C. Proteins were isolated from RPE/choroid as well as from retinal tissue of laser-treated and control mice as described above (ELISA). Protein concentration was evaluated by a colorimetric assay (PierceTM BCA Protein Assay Kit, 23225, Thermo Fisher Scientific). A multiarray electrochemiluminescence panel (U-Plex, Meso Scale Discovery, Rockville, Maryland, USA) was used to determine protein concentrations in RPE/choroid and retinal samples according to the manufacturer’s instructions. The following proteins were simultaneously measured with the kit: interleukin (IL) 6, IL-33, C-X-C motif chemokine ligand (CXCL) 2, CXCL10, CC-chemokine ligand (CCL) 2, CCL5, matrix metallopeptidase 9 (MMP9), tumor necrosis factor-alpha (TNFa) and Vascular Endothelial Growth Factor A (VEGFA).

### Statistics

Statistical analysis was performed using RStudio (Version 1.2.1335, R Version 3.5.3). Data were tested for normality applying the Kolmogorov–Smirnov test. If normality was given, an unpaired t-test or one-way ANOVA was applied, if not indicated otherwise. If the data did not meet the criteria of normality, the Mann–Whitney U test was applied. To compare CNV sizes between two groups, a paired Mann-Whitney U test was applied. Differences were considered significant when p value < 0.05.

### Study approval

For human samples, ethics approval was granted from local Ethics Committees. Written informed consent was received from participants prior to inclusion in the study. Participants were identified by number, not by name. All animal experiments were approved by the local authority (Regierungspräsidium Freiburg, Germany) and were performed in accordance with the respective national, federal and institutional regulations.

## Results

### Transcriptional characterization of human and mouse CNV

Transcriptional profiling identified 3043 and 2379 differentially expressed genes (DEG) in human and mouse CNV, when compared to the respective control tissue (Fig. 1A, B). An orthologous gene of the other species was found in the majority of all DEG (n = 4305, 87.4 %), whereas no orthologous gene existed for 606 (12.3 %) mouse and 15 (0.3 %) human DEG (Fig. 1B). Among the orthologous genes, about half were regulated in the same direction in both species, whereas the other half was expressed inversely (Fig. 1B).

A closer look at the differentially expressed genes revealed that several known CNV-associated factors were among the upregulated genes in both species, such as *MMP9* (Matrix Metallopeptidase 9), *PDGFB* (Platelet Derived Growth Factor Subunit B) and *AIF1* (Allograft Inflammatory Factor 1, IBA1) in human CNV (Fig. 2A) and *Cx3cr1* (C-X3-C Motif Chemokine Receptor 1), *Iqgap1* (IQ Motif Containing GTPase Activating Protein 1) and *Aif1* in murine CNV (Fig. 2B). In addition, *S100A8* (S100 calcium binding protein A8), *S100A9* and *MALAT1* (Metastasis Associated Lung Adenocarcinoma Transcript 1) were among the five most upregulated genes in human CNV (Fig. 2A), while *Gnat1* (G Protein Subunit Alpha Transducin 1), *Sag* (S-Antigen Visual Arrestin) and *Slc17a7* (Solute Carrier Family 17 Member 7) were among the five most upregulated genes in mouse CNV (Fig. 2B). Gene ontology (GO) analysis revealed that upregulated genes in human CNV were mainly involved in biological processes such as blood vessel development, response to wounding, actin cytoskeleton organization and immune processes like myeloid leukocyte activation and cytokine production (Fig. 2C). In mouse CNV membranes, upregulated genes were pre-dominantly associated to synapse organization, cytokine production, actin cytoskeleton organization and blood vessel development (Fig. 2E). Looking at the proportion of CNV specific genes associated to a disease-relevant biological process, a higher percentage of upregulated genes in human CNV was involved in angiogenesis (8.4 %) and wounding (6.9 %) compared to mouse CNV (4.8 % and 0.5 %, respectively). In contrast, mouse CNV was characterized by a higher proportion of immune-related genes (10.2 % vs. 4.4 %, Fig. 2D). Analyzing the factors involved in blood vessel development in both species, the top five expressed genes were *APOE* (apolipoprotein E), *COL1A1* (collagen type I alpha 1 chain), *COL1A2* (collagen type I alpha 2 chain), *GPX1* (glutathione peroxidase 1) and *SPARC* (secreted protein acidic and cysteine rich) in human CNV (Fig. 2F) and *Hspg2* (heparan sulfate proteoglycan 2), *Mfge8* (milk fat globule-EGF factor 8 protein), *Tnfrsf1a* (tumor necrosis factor receptor superfamily, member 1a), *Adam15* (a disintegrin and metallopeptidase domain 15) and *Col4a2* (collagen, type IV, alpha 2) in murine CNV (Fig. 2G). Among the genes associated to cytokine production, *S100A9, CD74, CYBA* (cytochrome b-245 alpha chain), *FN1* (fibronectin 1) and *RPS3* (ribosomal protein S3) were the five most highly expressed genes in human CNV (Fig. 2F) and *Iqgap1, Ptprs* (protein tyrosine phosphatase, receptor type, S), *Mertk* (MER proto-oncogene tyrosine kinase), *Mapk3* (mitogen-activated protein kinase 3) and *Sirpa* (signal-regulatory protein alpha) in murine CNV (Fig. 2G).

**Figure 2:**
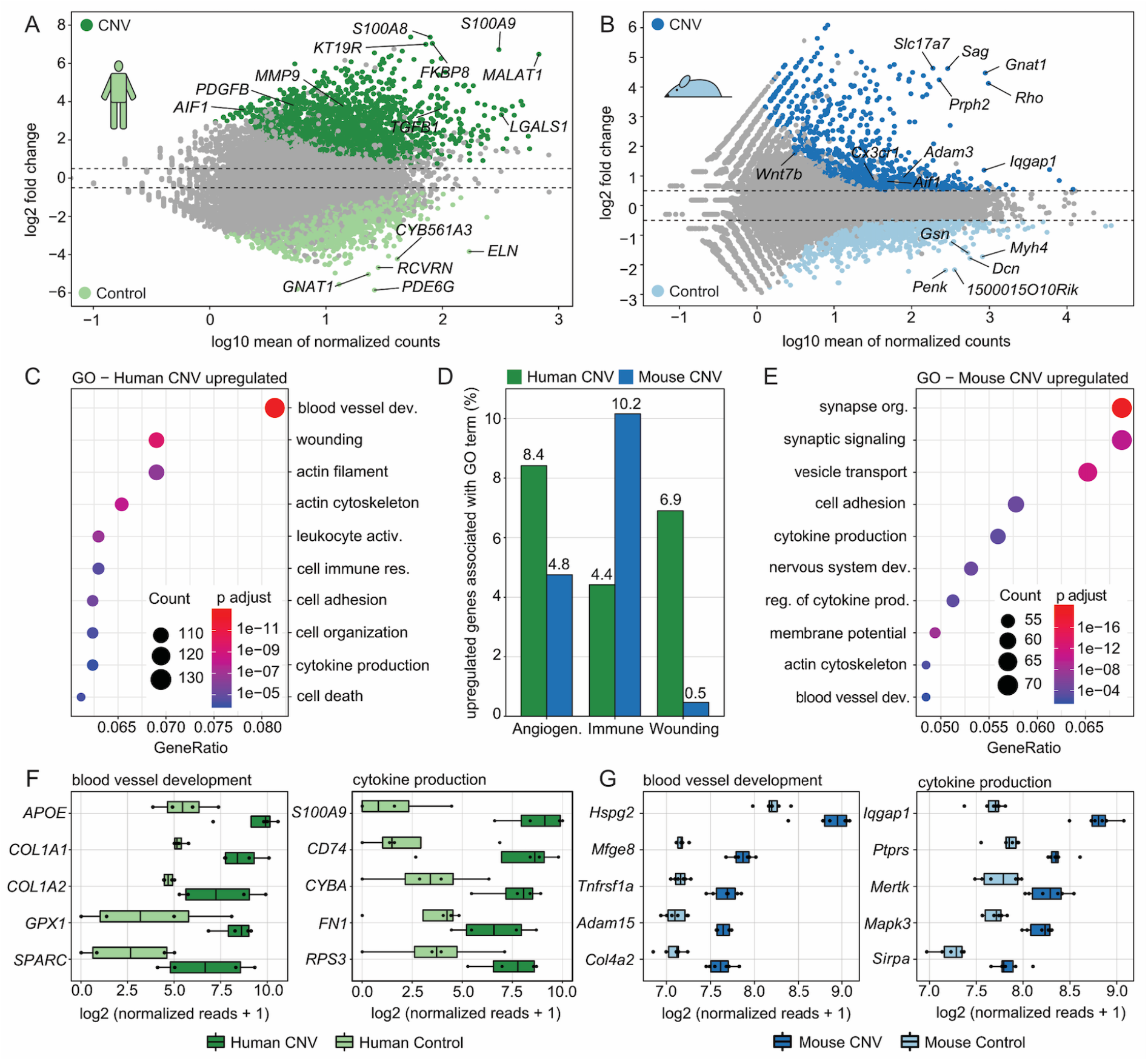
Transcriptional characterization of human and mouse CNV. (A-B) MA plots illustrating differentially expressed genes (DEG) between (A) human CNV (dark green, log2FC > 0.5 and adjusted p < 0.05, n = 1766) and human control tissue (light green, log2FC < -0.5 and adjusted p < 0.05, n = 1277) as well as between (B) mouse CNV (dark blue, log2FC > 0.5 and adjusted p < 0.05, n = 1196) and mouse control (light blue, log2FC < - 0.5 and adjusted p < 0.05, n = 1183). The top five factors according to the product of log2FC and mean of normalized reads as well as five relevant CNV-associated DEGs known from the literature are labeled. (C+E): Dotplots showing the top ten Gene ontology (GO) biological processes, which the significantly upregulated genes in human (C) and mouse (E) CNV were involved in. The size of the dots represents the number of associated genes (count). The adjusted p-value of each GO term is shown by color. The gene ratio describes the ratio of the count to the number of all DEGs. (D): Barplot visualizing the percentage of upregulated genes involved in three disease-relevant biological processes for human and mouse CNV: angiogenesis (GO:0001525, GO:0001568), immune response (GO:0006955) and response to wounding (GO:0009611). (F-G): Box plots illustrating the normalized reads of the top five expressed genes of two disease-relevant GO terms (GO:0001568, GO:0001816) according to mean expression in CNV samples for human (F) and mouse (G).

### Phylogenetically conserved mediators of CNV

To identify conserved molecular patterns between mice and men, we first determined genes which were significantly upregulated in CNV membranes of both species when compared to the respective control tissue (Fig. 3A). The commonly upregulated genes in human and mouse CNV (n = 95) as well as species-specific DEG (mouse: n = 151, human: n = 802) are illustrated in Figure 3A. GO analysis revealed that the 95 shared genes were predominantly involved in biological processes such as blood vessel development/angiogenesis, immune response signal transduction, apoptotic signaling pathway, myeloid leukocyte activation, leukocyte degranulation and response to wounding (Fig. 3B). The genes involved in these biological processes are visualized in Figure 3C. Following this approach, genes such as *FN14* (fibroblast growth factor inducible-14), *LGALS3* (galectin 3), *AIF1, CTSS* (cathepsin S), *UNC5B* (unc-5 netrin receptor B), *ADAM15, MCAM* (melanoma cell adhesion molecule), *CYBB* (cytochrome b-245 beta chain), *APLNR* (apelin receptor), *KCNN4* (potassium calcium-activated channel subfamily N member 4) and *UNC93B1* (unc-93 homolog B1, TLR signaling regulator) were identified as conserved mediators of CNV across both species (Fig. 3A and 3C). Since transcriptional profiling identified *AIF1* (also known as IBA1) as one of the phylogenetically conserved mediators in human and mouse CNV, immunohistochemical validation was subsequently performed. These experiments confirmed significantly increased accumulation of AIF1-positive cells in the area of CNV in human (Supplement Fig. 1A) and mouse CNV (Supplement Fig. 1C) when compared to the respective control tissue (Supplement Fig. 1B and 1D).

**Figure 3:**
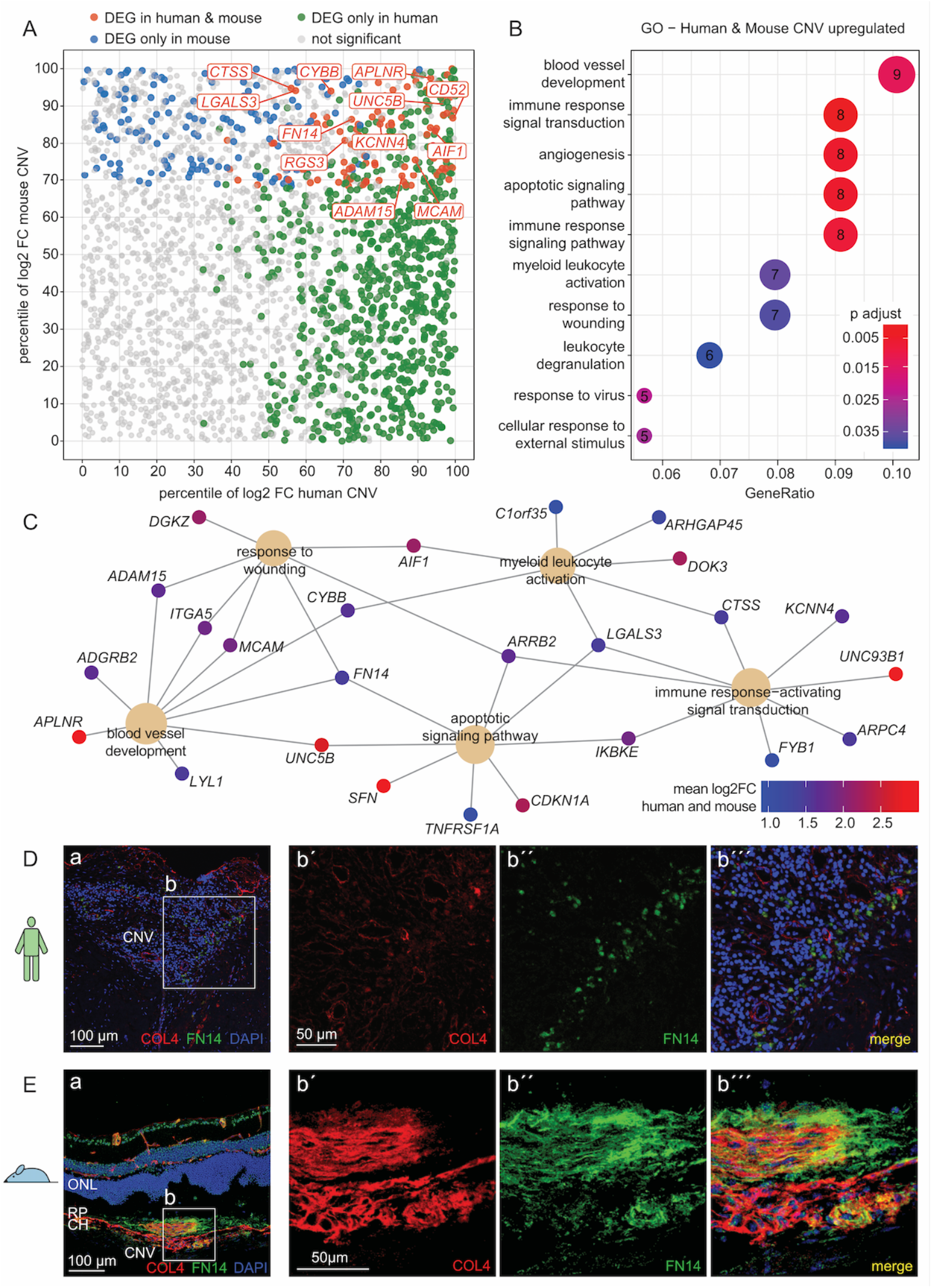
Phylogenetically conserved mediators of murine and human CNV. (A): Scatter plot illustrating genes significantly upregulated in both human and mouse CNV (n = 95, red), only in human CNV (n = 802, green) or only in mouse CNV (n = 151, blue) (definition: log2FC > 0.5 and adjusted p < 0.05). The percentile of log2 fold change between CNV and control samples for human (x-axis) as well as for mouse (y-axis) was calculated for every gene. (B): Dotplot showing the top ten Gene ontology (GO) biological processes, which the 95 significantly upregulated genes in both species were involved in (red in A). The number of associated genes is indicated within each circle. The adjusted p-value of each GO term is color-coded. The gene ratio describes the ratio of the count to the number of all DEG (n = 95). (C): Cnetplot illustrating phylogenetically conserved genes (small circles) associated with the top biological processes (large circles) from (B). Color indicates mean log2 fold change of human and mouse CNV compared to the respective control group. FN14 was identified as a key factor among the patho-physiologically most relevant biological processes, with a 4.1-fold increase in human and 1.8-fold increase in murine CNV. (D-E): Immunohistochemical validation of FN14 in human CNV (D) as well as in murine laser-induced CNV (E). (D): a: section of human CNV membrane reveals significant immunoreactivity against FN14. b: higher magnification images of FN14 expression in human CNV. Vessels are stained for COL4 (in red) and nuclei are counterstained with DAPI (in blue). (E): a: cryosection of murine laser-induced CNV demonstrating high FN14 immunoreactivity in CNV as well as some expression in physiological retinal vasculature and nuclei, especially in ganglion cell and inner nuclei layers. b: higher magnification images of FN14 expression in CNV. Vessels are stained for COL4 (in red) and nuclei are counterstained with DAPI (in blue). Abbreviations: CH: choroid, CNV: choroidal neovascularization, ONL: outer nuclear layer, RP: retinal pigment epithelium.

### FN14 and its ligand TWEAK in human and mouse CNV

RNA sequencing identified FN14 as a conserved mediator in human and murine CNV representing a key factor among the pathophysiologically most relevant biological processes (Fig. 3C). To validate these findings on a protein level, the expression of FN14 and its localization in human and murine CNV as well as control tissue were further investigated by immunohistochemistry. In line with the RNA sequencing results, an increased immunoreactivity against FN14 was observed in human and mouse CNV (Fig. 3D and 3E) when compared to control tissue (Supplement Fig. 2A and 2B). Additionally, there was a less pronounced but still detectable immunoreactivity in physiological murine choroidal and retinal vessels as well as within the inner nuclear layer and ganglion cell layer which was not distinctive between CNV and control samples (Fig. 3E and Supplement Fig. 2B). Negative controls without primary antibody against FN14 as well as isotype control revealed a high specificity of the same primary antibody used in human and mouse samples as well as secondary antibodies (Supplement Fig. 3). In addition, the concentration of the FN14 ligand TWEAK (tumor necrosis factor-like weak inducer of apoptosis) (Ameri et al., 2014) was measured by ELISA. TWEAK was detectable in all human vitreous samples as well as in the plasma of control and nAMD patients (Supplement Fig. 4A). In both the control and nAMD groups, vitreous concentrations were 2.6 and 3.0 times higher than in plasma, respectively, indicating a local accumulation. The TWEAK concentration was comparable between controls and CNV in vitreous (585.0 ± 68.2 pg/ml vs. 498.5 ± 54.7 pg/ml) and plasma samples (222.6 ± 92.6 pg/ml vs. 164.0 ± 33.7 pg/ml) (Supplement Fig. 4A). In line with the results in human, TWEAK was detected in all murine samples with comparable concentrations between the CNV and the CNV group in RPE (139.2 ± 7.8 pg/ml vs. 147.9 ± 7.5 pg/ml) and retinal tissue (80.5 ± 4.0 pg/ml vs. 72.7 ± 4.7 pg/ml) (Supplement Fig. 4B).

**Figure 4:**
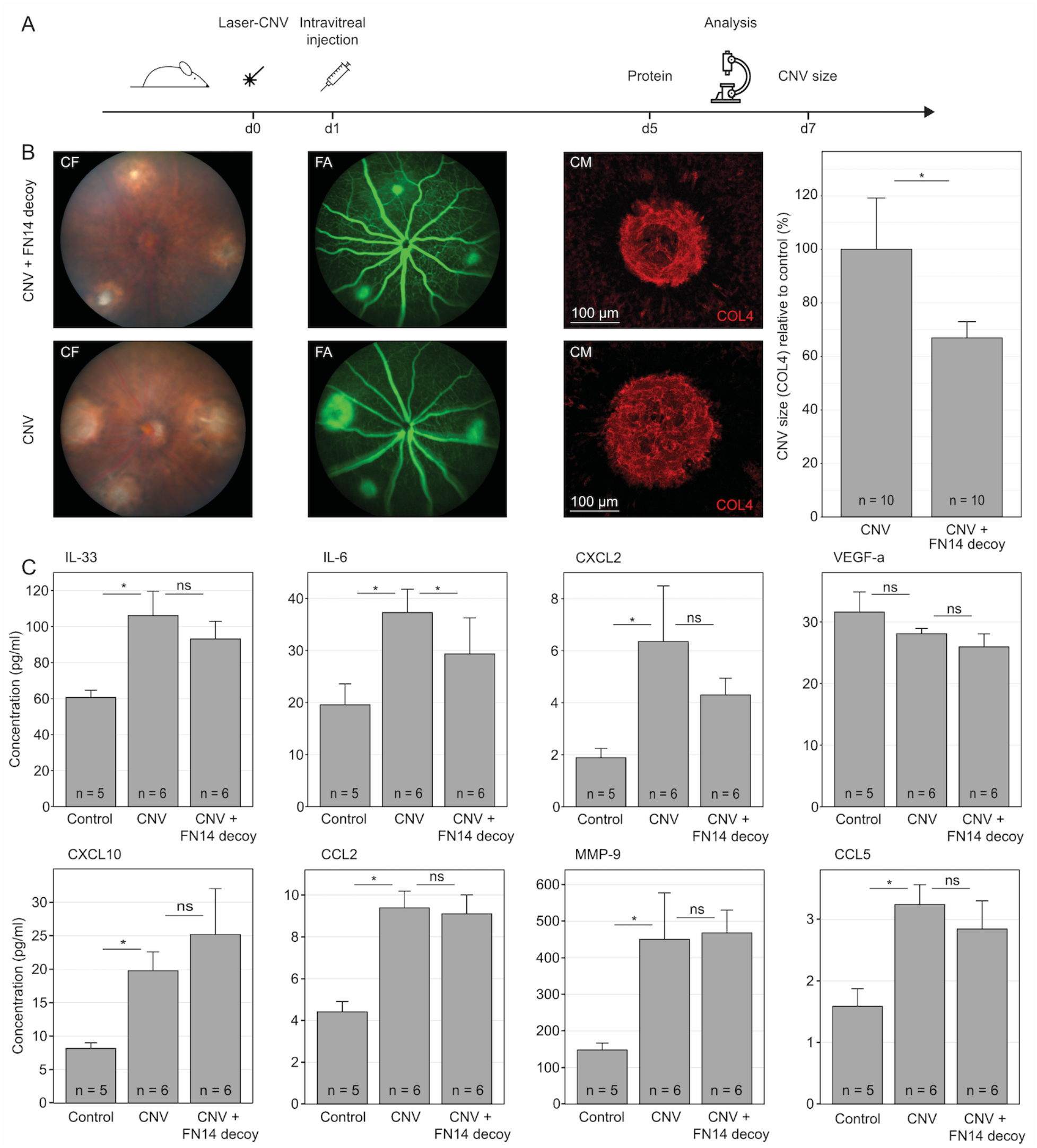
FN14 decoy receptor modulates the cytokine profile in CNV and reduces neovascularization size. (A): Experimental setup. (B): Intravitreal injection of a FN14 decoy receptor compared to PBS (control) significantly reduced CNV size. Color fundus (CF) and fluorescence angiography (FA) on d7 of representative eyes of the FN14 decoy receptor (upper panel) and the PBS group (lower panel) as well as confocal microscopy (CM) images of representative lesions (COL4 in red) the size of which approximately corresponds to the mean spot size within the respective group. (C): Protein concentrations of several cytokines on d5 in the RPE of unlasered controls, CNV injected with PBS (CNV) or with FN14 decoy receptor on d1 (CNV + decoy). Data are shown as mean with SEM. For statistical analysis, paired ANOVA adjusting for total protein concentration was applied. *: p<0.05, ns: not significant.

### FN14 decoy receptor modulates the cytokine profile in CNV and reduces neovascularization

The substantial upregulation of FN14 RNA and protein in human and murine CNV suggests a conserved role in the pathophysiology of CNV formation. To further investigate this hypothesis, a FN14 decoy receptor, which blocks the interaction between FN14 and its ligand TWEAK, was injected intravitreally on day 1 after laser CNV induction in 10 murine eyes. PBS injected into the partner eye served as control (Fig. 4A). Interestingly, intravitreal injection of FN14 decoy receptor resulted in a significant reduction (33.1 %, SEM: 6.0, p<0.05) of CNV size compared to PBS injected eyes (Fig. 4B).

To investigate the effect of the FN14 decoy receptor on the cytokine milieu in CNV, eight disease-relevant cytokines were measured in CNV tissue at d5 following FN14 decoy receptor treatment at d1. PBS injected into each contralateral eye as well as 5 unlasered eyes served as controls (Fig. 4C). CNV induction was associated with a significant increase in various cytokines on day 5, among them IL-33, IL-6, CXCL2, CCL2, MMP-9 and CCL5 (Fig. 4C). Interestingly, in CNV eyes, the FN14 decoy receptor led to a significant reduction of IL-6 when compared to PBS-injected eyes (29.3 ± 6.9 pg/ml vs. 37.3 ± 4.5 pg/ml, p<0.05, Fig. 4C). IL-33, CXCL2 and CCL5 also tended to be decreased in the FN14 decoy group, although the differences were not significant (Fig. 4C). Of note, inhibition of FN14 did not significantly affect VEGF-a concentration on day 5 after CNV induction (Fig. 4C), suggesting a VEGF-independent anti-angiogenic effect of the FN14 inhibition.

## Discussion

Anti-VEGF therapy has significantly improved visual outcome of patients with neovascular AMD during the last years (Bressler et al., 2011; Mitchell et al., 2018). However, about one third of patients show persistent exudation and a slow decrease in visual acuity despite recurrent anti-VEGF injections (Wecker et al., 2019; Yang et al., 2016). This suggests additional mediators that lead to differentiated vascular membranes or subretinal fibrosis in the final stage of the disease. The present study performed comparative transcriptomics between human and murine CNV in order to identify cross species mediators of CNV formation. Following this approach, the study identifies fibroblast growth factor inducible-14 (FN14) as a so far unrecognized conserved factor in CNV formation, the inhibition of which results in reduced IL-6 expression and a reduction of CNV size *in vivo*.

The mouse model of laser-induced CNV has been used extensively in preclinical studies of nAMD as it mimics the hallmarks and complex sequelae of nAMD including rupture of Bruch’s membrane, immune cell activation, and subsequent acute formation of new choroidal blood vessels (Lambert et al., 2013). Our analysis shows that the transcriptional profile of both human and murine CNV is characterized by enhanced activation of biological processes such as blood vessel development, actin cytoskeleton organization, and cytokine production. However, CNV of both species differed by the predominance of angiogenesis and wound healing processes in human CNV and more pronounced immune processes in murine CNV, which reflects the more chronic nature of human CNV and the acute and traumatic etiology of CNV in the mouse model (Lambert et al., 2013). Nevertheless, parallels in the expression pattern of myeloid cells between mice and humans have been found during CNV formation in previous studies (Beguier et al., 2020; Schlecht et al., 2020b; Wieghofer et al., 2021). Of note, about half of the CNV-associated genes were inversely expressed between human and mouse, indicating distinct differences between the human disease and the mouse model on the transcriptional level. Despite a high number of species-specific factors, 95 DEG were identified that were significantly upregulated in both human and murine CNV when compared to the respective control tissue. These genes are known mediators of disease-relevant processes such as angiogenesis, immune response and wounding. Among them, there were several proangiogenic factors including *APLNR* (Hara et al., 2013; Ishimaru et al., 2017; Kasai, 2011; McAnally et al., 2018), *LGALS3* (Chen et al., 2017; Jia et al., 2013; Yao et al., 2019), *CYBB* (Al-Shabrawey et al., 2005; Chan et al., 2013), *CTSS* (Chen et al., 2010; Fan et al., 2012; Wang et al., 2006), *UNC5B* (Lejmi et al., 2008), *MCAM* (Stalin et al., 2016), *ADAM15* (Horiuchi et al., 2003; Hou et al., 2015; Xie et al., 2008) and *KCNN4* (Grgic et al., 2005). In addition to the already known factors, the present study identifies a plethora of new conserved mediators of CNV development that represent potential therapeutic targets for neovascular AMD. Among these factors, FN14 emerged as a key factor among the most significantly activated and disease-relevant processes in human and mouse CNV. Immunohistochemistry confirmed an increased expression of FN14 protein in the area of human and murine CNV. Besides, a weaker but still distinct immunoreactivity against FN14 was observed in the murine inner nuclear layer and ganglion cell layer as well as in physiological retinal and choroidal vessels, which is in accordance with the literature (Ameri et al., 2014). The authors suggest a baseline expression of FN14 that increases significantly under pathological conditions such as the development of CNV (Ameri et al., 2014). The increased expression of FN14 under pathological conditions, which is assumed to be mediated by growth factors, such as FGF, PDGF and VEGF (Burkly et al., 2007), is well known in the course of various pathologies (Burkly et al., 2007) including the formation of retinal neovascularization (Ameri et al., 2014). In addition, the FN14 lig- and TWEAK was distinctly detectable in all human vitreous as well as murine RPE and retinal samples, further indicating a highly conserved (Burkly et al., 2007) and potential pathophysiological involvement of the FN14-TWEAK pathway in human and mouse CNV. Interestingly, we did not observe a significant difference in TWEAK concentration in the human vitreous as well as murine RPE and retina between CNV and control groups. These results imply that in CNV, the FN14-TWEAK pathway may be predominantly regulated by the expression of FN14 rather than by its ligand TWEAK, which is consistent with the observations for retinal neovascularization (Ameri et al., 2014) and various other pathologies which an increased expression of FN14 was reported for (Burkly et al., 2007). It is interesting to note that TWEAK concentrations were significantly higher in human vitreous compared to the corresponding plasma samples of the same patients indicating a distinctive role of the pathway within the ocular compartment. The FN14-TWEAK axis is known for its proinflammatory and proangiogenic properties involved in processes such as endothelial cell proliferation (Harada et al., 2002), wound healing and tissue regeneration, contributing to a variety of pathologies, including retinal neovascularization (Abu El-Asrar et al., 2015; Ameri et al., 2014; Sun et al., 2020), inflammatory diseases (Winkles, 2008) and cancer (Winkles, 2008). Besides, TNF Receptor Superfamily Member 10a (TNFRSF10A), belonging to the same TNF receptor superfamily as FN14 (TNFRSF12A), has recently been associated to AMD in a genome wide association study (Fritsche et al., 2016) and, in accordance with the localization of FN14 observed in the present study, was found to be expressed most significantly in vascular cells of the human retina, as determined by single cell RNA sequencing (Menon et al., 2019). However, the role of the FN14 signaling pathway has not yet been evaluated in CNV.

To further investigate the pathophysiological significance of the FN14-TWEAK pathway in CNV formation, a FN14 decoy receptor, which blocks the interaction between FN14 and its ligand TWEAK, was injected intravitreally, resulting in a significantly reduced CNV size *in vivo*. Similar results have been obtained for retinal neovascularization, where inhibition of the FN14-TWEAK signaling pathway resulted in attenuated vessel formation (Ameri et al., 2014). To gain further insights into the antiangiogenic mechanisms, the cytokine milieu in the CNV tissue was investigated *in vivo* by adding or omitting the FN14 decoy receptor. Blocking the FN14-TWEAK-pathway resulted in a significant reduction of IL-6 concentration in the CNV microenvironment with a tendency to also decrease IL-33, CXCL2 and CCL5, although the differences were not significant. These results are in accordance with studies on human astrocytes (Saas et al., 2000) and orbital fibroblasts (Lee et al., 2018) as well as murine synovial (Kamata et al., 2006) and mesangial cells (Campbell et al., 2006), which have shown FN14-TWEAK activation to induce IL-6, CXCL2 and CCL5 secretion, respectively. These results may indicate that FN14 influences CNV development, at least in part, through modulation of IL-6, although this does not exclude other soluble factors to be regulated by the treatment with the FN14 decoy receptor. In consequence, targeting FN14 may be a novel promising therapeutic strategy for the treatment of CNV. Of note, clinical studies evaluating antibody-based inhibitors of the FN14-TWEAK signaling pathway have demonstrated a high level of tolerability (Wisniacki et al., 2013) and a significant reduction of FN14-TWEAK signaling in humans (Meulendijks et al., 2016), further supporting a potential therapeutic applicability for neovascular AMD.

Taken together, this study characterizes the transcriptome of human and mouse CNV membranes in an unpreceded and unbiased manner and identifies FN14 as a so far unrecognized phylogenetically conserved mediator of CNV formation. Intravitreal injection of an FN14 decoy receptor significantly reduced IL-6 expression and CNV size *in vivo*. FN14 is thus suggested as a promising novel therapeutic target for neovascular AMD, which may potentially be beneficial in patients with exudative AMD.

## Acknowledgements

The authors thank Lutz Hansen for surgical assistance, Gabriele Prinz and Sylvia Zeitler for excellent technical assistance. This study was supported by the Helmut-Eckert foundation and Volker Homann foundation.

## Author contributions

JW: designing research studies, conducting experiments, acquiring data, analyzing RNA-sequencing data, writing the manuscript, AS: designing research studies, conducting experiments, review and editing the manuscript, DDR: conducting experiments, review and editing the manuscript, SB: review and editing the manuscript, HA: review and editing the manuscript, GS: supervising experiments, review and editing the manuscript, PW: conducting experiments, supervising experiments, review and editing the manuscript, CL: designing research studies, analyzing RNA-sequencing data, supervising experiments, review and editing the manuscript.

---

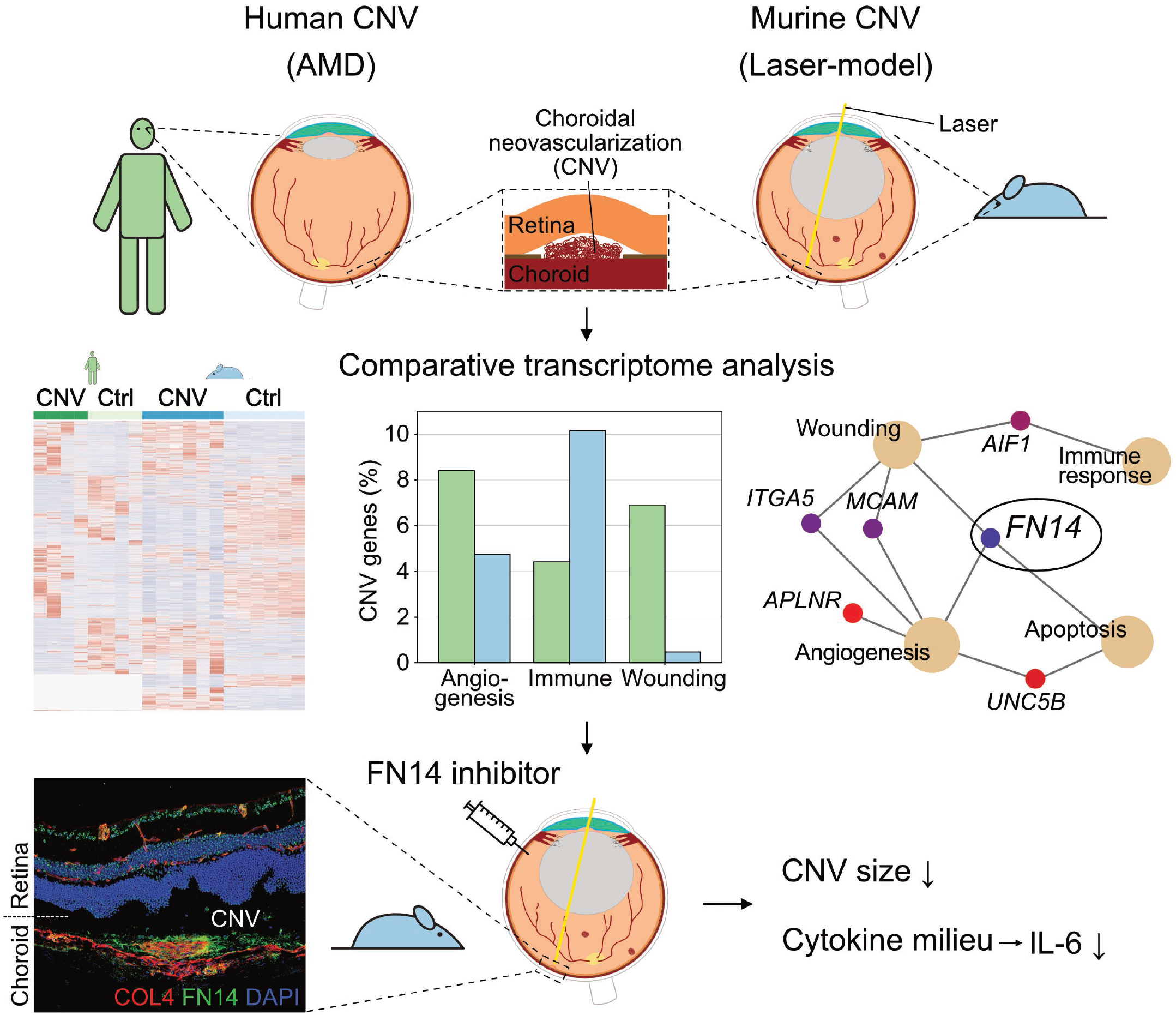

**Supplementary Figure 1:**
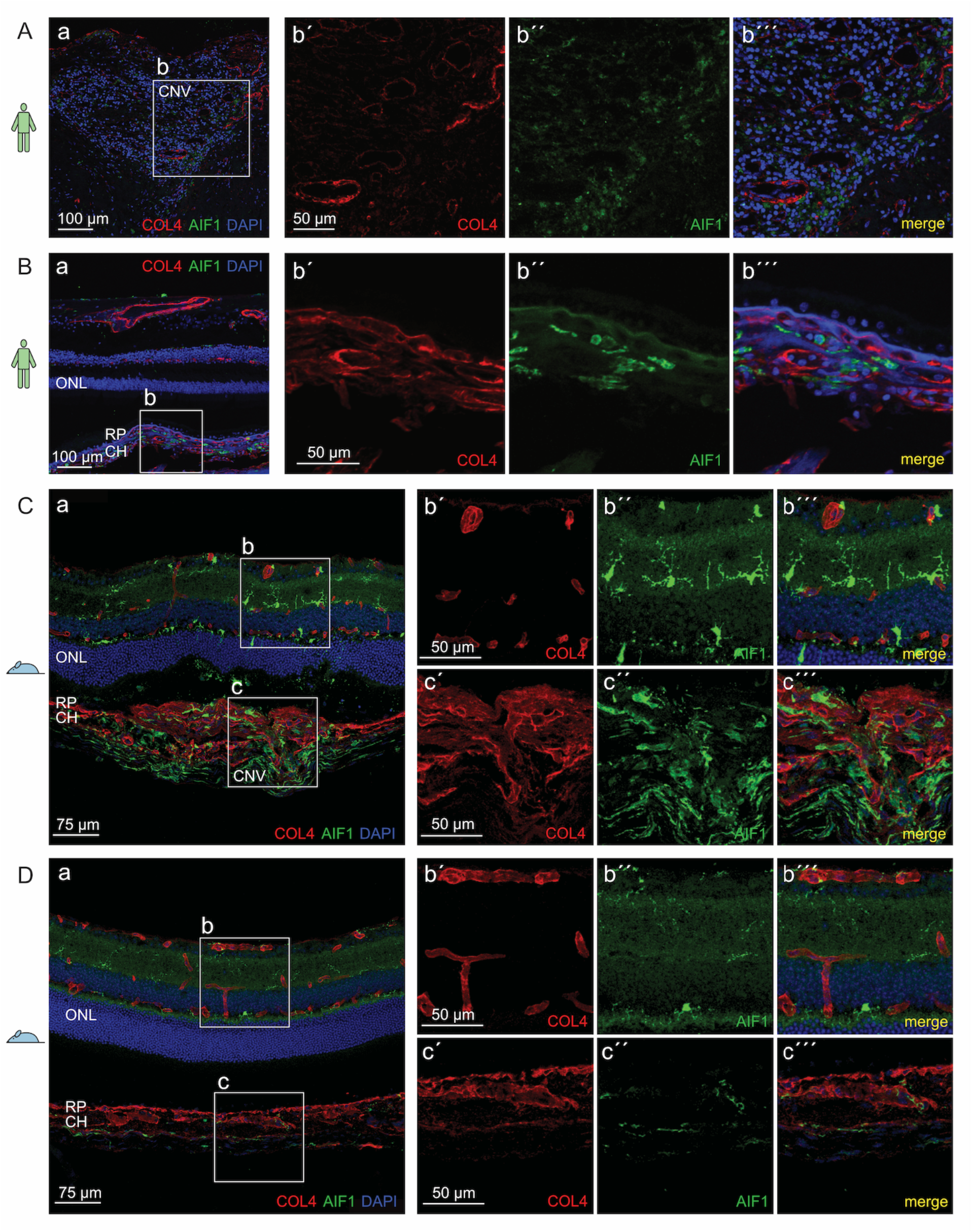
Immunohistochemical validation of AIF1 in human (A-B) and mouse (C-D) CNV (A+C) and control tissue (B+D). (A): a: sections of human CNV membranes demonstrating accumulation of AIF1 positive cells (in green) at sites of CNV stained for COL4 (in red). b: higher magnification images of AIF1-positive cells in the area of CNV. Vessels are stained for COL4 (in red) and nuclei are counterstained with DAPI (in blue). (B): control tissue shows significantly lower number of AIF1-positive cells when compared to CNV. (C): a: cryosections of mouse laser-induced CNV demonstrating accumulation of AIF1-positive cells (in green) in the area of CNV stained for COL4 (in red). Right panel: higher magnification images of AIF1-positive cells in the retina (upper panel) as well as in the area of CNV (lower panel). Vessels are stained for COL4 (in red) and nuclei are counterstained with DAPI (in blue). (D): significantly lower number of AIF1-positive cells in murine control tissue. Abbreviations: CH: choroid, CNV: choroidal neovascularization, ONL: outer nuclear layer, RP: retinal pigment epithelium.

**Supplementary Figure 2:**
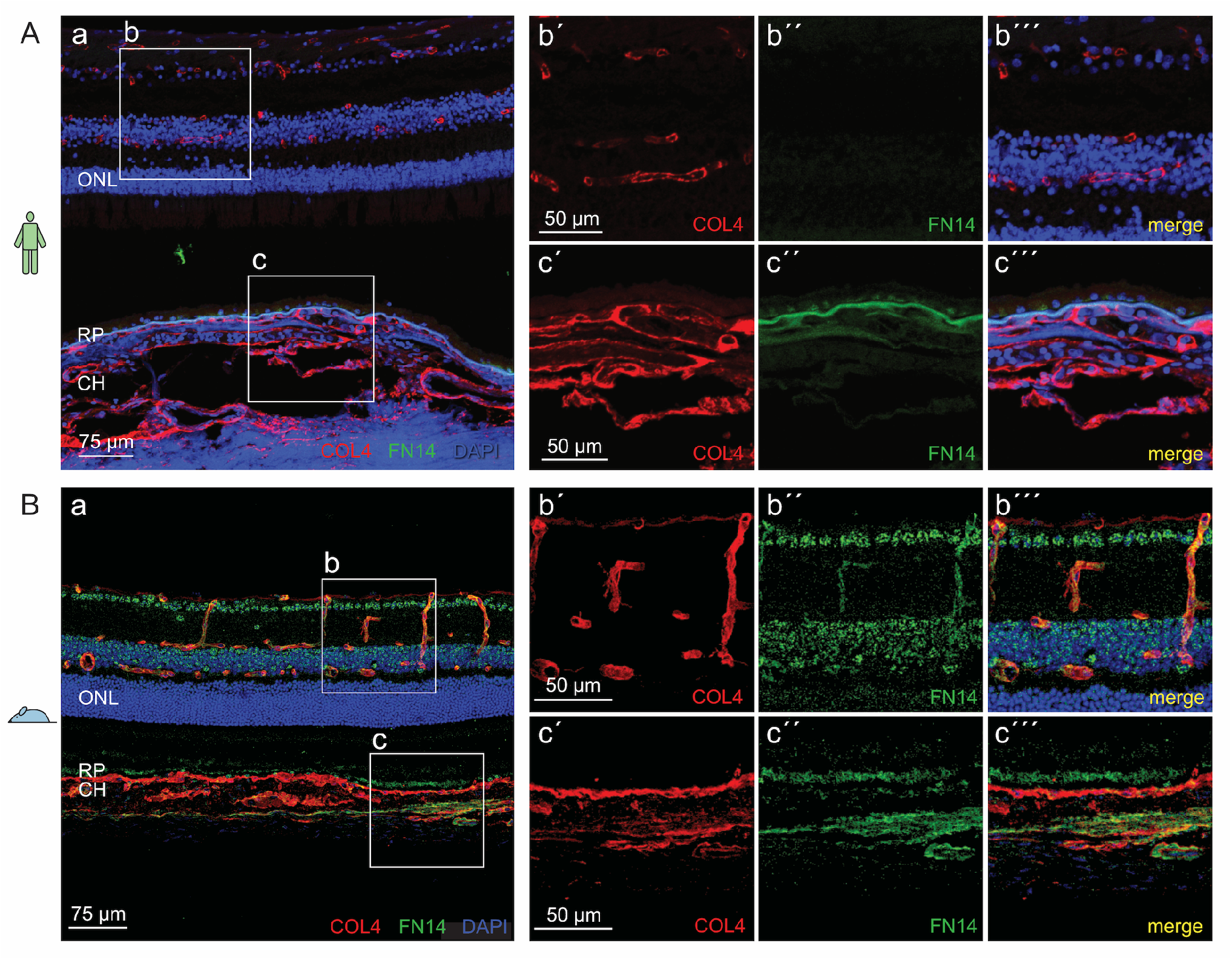
FN14 staining in human (A) and mouse (B) control tissue revealing significantly lower immunoreactivity in RPE/choroid (c) compared to CNV (Fig. 3D and E). Vessels and CNV are stained for COL4 (in red) and nuclei are counterstained with DAPI (in blue). Abbreviations: CH: choroid, ONL: outer nuclear layer, RP: retinal pigment epithelium. The areas within the white continuous line are shown in higher magnification to the right.

**Supplementary Figure 3:**
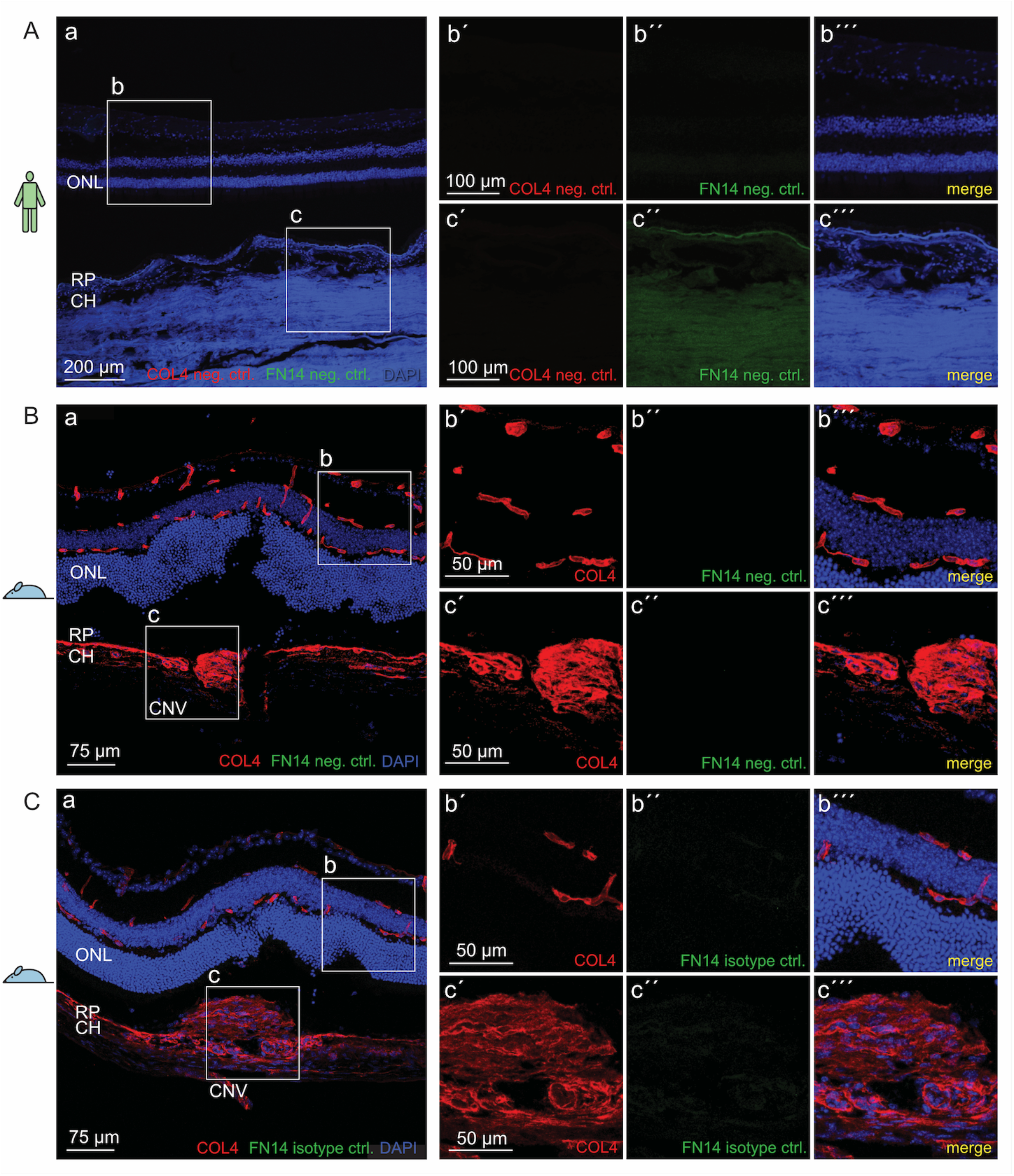
(A+B) FN14 negative control as well as (C) FN14 isotype control in human CNV (A) as well as in laser-induced CNV (B-C) revealed high specificity of primary and secondary antibodies. Vessels are stained for COL4 (in red) and nuclei are counterstained with DAPI (in blue). Abbreviations: CH: choroid, CNV: choroidal neovascularization, ONL: outer nuclear layer, RP: retinal pigment epithelium.

**Supplementary Figure 4:**
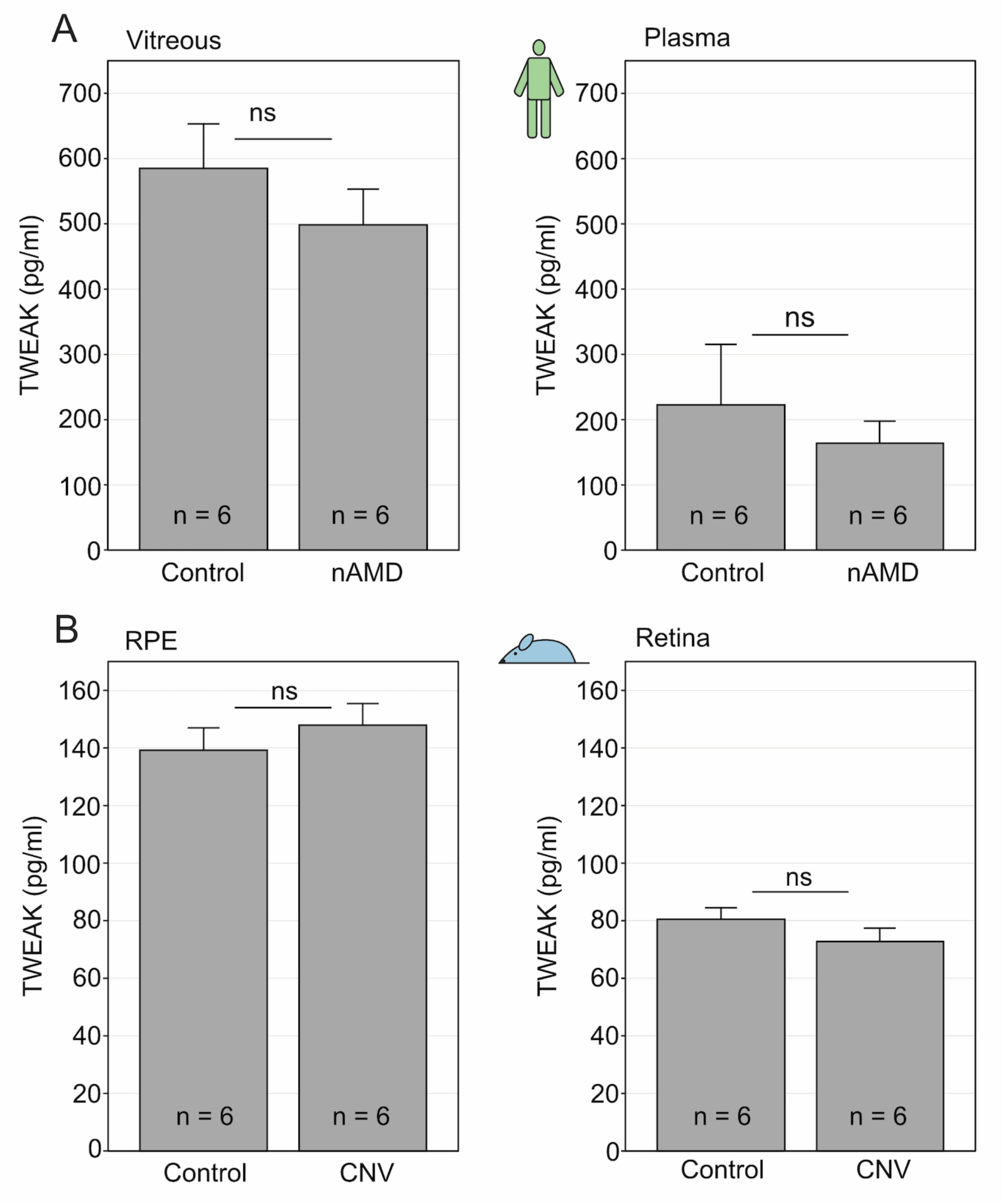
(A): Vitreous and plasma concentration of TWEAK in neovascular AMD and control patients as well as in (B) murine RPE and retina samples in CNV and control group as determined by TWEAK ELISA. Data is shown as mean with SEM. Plasma samples in humans were taken at the time of surgery.

